# Peripheral blood DNA methylation and autism spectrum disorder

**DOI:** 10.1101/320622

**Authors:** Shan V. Andrews, Brooke Sheppard, Gayle C. Windham, Laura A. Schieve, Diana E. Schendel, Lisa A. Croen, Pankaj Chopra, Reid S. Alisch, Craig J. Newschaffer, Stephen T. Warren, Andrew P. Feinberg, M. Daniele Fallin, Christine Ladd-Acosta

## Abstract

Background

Several reports have suggested a role for epigenetic mechanisms in ASD etiology. Epigenome-wide association studies (EWAS) in autism spectrum disorder (ASD) may shed light on particular biological mechanisms. However, studies of ASD cases versus controls have been limited by post-mortem timing and severely small sample sizes. Reports from in-life sampling of blood or saliva have also been very limited in sample size, and/or genomic coverage. We present the largest case-control EWAS for ASD to date, combining data from population-based case-control and case-sibling pair studies.

Methods

DNA from 968 blood samples from children in the Study to Explore Early Development (SEED 1) was used to generate epigenome-wide array DNA methylation (DNAm) data at 485,512 CpG sites for 453 cases and 515 controls, using the Illumina 450K Beadchip. The Simons Simplex Collection (SSC) provided 450K array DNAm data on an additional 343 cases and their unaffected siblings. We performed EWAS meta-analysis across results from the two data sets, with adjustment for sex and surrogate variables that reflect major sources of biological variation and technical confounding such as cell type, batch, and ancestry. We compared top EWAS results to those from a previous brain-based analysis. We also tested for enrichment of ASD EWAS CpGs for being targets of meQTL associations using available SNP genotype data in the SEED sample.

Findings

In this meta-analysis of blood-based DNA from 796 cases and 858 controls, no single CpG met a Bonferroni discovery threshold of p < 1.12×10^−7^. Seven CpGs showed differences at p < 1×10^−5^ and 48 at 1×10^−4^. Of the top 7, 5 showed brain-based ASD associations as well, often with larger effect sizes, and the top 48 overall showed modest concordance (r = 0.31) in direction of effect with cerebellum samples. Finally, we observed suggestive evidence for enrichment of CpG sites controlled by SNPs (meQTL targets) among the EWAS CpGs hits, which was consistent across EWAS and meQTL discovery p-value thresholds.

Conclusions

We report the largest case-control EWAS study of ASD to date. No single CpG site showed a large enough DNAm difference between cases and controls to achieve epigenome-wide significance in this sample size. However, our results suggest the potential to observe disease associations from blood-based samples. Among the 7 sites achieving suggestive statistical significance, we observed consistent, and stronger, effects at the same sites among brain samples. Discovery-oriented EWAS for ASD using blood samples will likely need even larger samples and unified genetic data to further understand DNAm differences in ASD.

## FINDINGS

The etiology of autism spectrum disorder (ASD) may involve epigenetic mechanisms. Indirect evidence supporting this hypothesis comes from the observation that children with Rett, Fragile X, and Angleman Syndromes often show impaired communication and exhibit repetitive behaviors [1–3], two core domains affected in autism. All 3 of these Syndromes are caused by epigenetic defects [4–9]. Additional evidence stems from genetic studies of rare variation in non-syndromic forms of ASD. Although these studies have primarily identified private variants associated with ASD, there is now strong evidence that the ASD associated variants converge upon three biological pathways, one of which is chromatin remodeling [10–12]. Finally, there is direct evidence from case-control postmortem brain studies supporting epigenetic involvement in ASD. Several candidate gene-based studies have shown altered epigenetic states associated with autism [13–18]. Genome-scale screens have identified changes in DNA methylation (DNAm) at specific CpG sites [19, 20] as well as global changes in non-CpG methylation levels [21] in postmortem cerebral tissue from individuals with ASD relative to controls. Studies of cerebral cortex tissue has revealed genomic spreading of histone H3 lysine 4 methylation and histone H3 lysine 27 acetylation marks, away from the promoter region, among a subset of individuals with ASD compared to controls [22, 23].

Examination of the affected tissue, i.e. brain, can provide important insights into potential mechanisms of disease etiology; however, there are considerable limitations with these types of studies. They suffer from severely small sample sizes, have historically had low genomic coverage, and often lack comprehensive unified clinical, demographic, and genomic data. Importantly, they are based on autopsy-derived tissue and do not reflect epigenetic marks in a living individual, are not at optimal developmental timing, and may be influenced by life experiences and cause of death. To overcome these barriers, complementary, large population-based autism epigenetic studies using accessible tissues, such as blood, from living individuals are needed. To date, three genome-scale epigenetic studies of autism in accessible peripheral tissues have been performed. One study of peripheral blood from 50 monozygotic twin pairs, including 6 pairs discordant for ASD at age 15, examined DNAm at over 27,000 CpG sites in promoter regions. The authors found suggestive evidence for epigenetic alterations associated with ASD and associated traits within families [24]. Similarly, an investigation of DNAm at CpG island regions in lymphoblastoid cell lines, obtained from 7 twin pairs including 3 discordant for ASD, found ASD-related DNAm changes at the *RORA* gene [25]. Both of these studies were limited by the small number of samples examined, lack of genome-scale coverage, and specific focus on twin pairs with a lack of extension to the general population. Ectoderm cell lineage derived buccal cells, obtained from 47 ASD cases and 48 controls born to mothers aged 35 and older, have also shown suggestive epigenetic alterations associated with ASD [26]. While suggestive, it is unclear how these buccal-based epigenetic findings relate to a population sample and in a larger number of individuals. Thus, more research in accessible tissues from larger population-based, non-familial, samples is needed.

Here, we overcome previous limitations and perform the largest epigenome-scale examination of DNAm, to date, among two large U.S. case-control studies of autism: the Study to Explore Early Development, phase I (SEED I) and the Simons Simplex Collection (SSC). Both measured DNAm at over 450,000 loci in childhood blood samples from either population-based cases and controls (SEED I) or discordant sibling pairs (SSC). Meta-analysis across both sets included 796 ASD cases and 858 controls. In addition to CpG-specific differential DNAm, we explored the set of blood-derived differentially methylated sites for their concordance in post-mortem brain tissue and their enrichment for genetically-controlled CpG sites.

## Methods

### Study to Explore Early Development (SEED)

The Study to Explore Early Development is a multi-site case-control study with population-based ascertainment. In SEED Phase 1, a total of 3,899 families were recruited across 6 study sites (California, Colorado, Georgia, Maryland, North Carolina and Pennsylvania) and classified into 3 groups according to child’s diagnosis: an autism spectrum disorder (ASD) group, a general population control group (POP), and a (non-ASD) developmental delay group. Details regarding participant recruitment, biospecimen collection, and final outcome classification have been previously described [27, 28]. Briefly, eligible children were born in one of the catchment areas between September 1, 2003 and August 31, 2006, which corresponded to being aged 2-5 years at the time of SEED phase I enrollment, resided in the same catchment area at the time of initial contact, and were required to live with a knowledgeable caregiver who could communicate in English (or in English or Spanish in California or Colorado) [27]. Biospecimens were collected when the children were between the ages of 3 and 5 years. Children with possible ASD and DDs were ascertained through multiple sources providing services for children with developmental disorders including hospitals, individual providers, clinics, and education and intervention programs. Parents with a child with an ASD or DD diagnosis could also contact the study directly to enroll. General population controls were ascertained through random sampling of vital records in the catchment areas [27]. This provides a more diverse segment of the population than solely recruiting participants from autism clinics.

Primary caregivers completed the Social Communications Questionnaire (SCQ) [29], a screener for autism spectrum disorder, during the study invitation phone call. Children with an SCQ score below 11 and without a previous ASD diagnosis were asked to participate in a general developmental evaluation in the clinic using the Mullen Scale of Early Learning (MSEL) [30]. If the SCQ score was above 11, the child had previously received an ASD diagnosis, or a clinician suspected ASD during the clinic visit, the child additionally completed a full ASD evaluation that included the Autism Diagnostic Observation Schedule (ADOS) [31–33] and the Autism Diagnostic Interview Revised (ADI-R) [34, 35]. ASD was confirmed based on scores on the ADI-R and ADOS, as described in detail elsewhere [36]. Institutional review boards at each study site and at the Centers for Disease Control and Prevention (CDC) approved the SEED study. Informed consent was obtained from all enrolled participants. For this study, we measured methylation among a subset of SEED Phase 1 individuals (n=980) with genome-wide genotyping data, a complete caregiver interview, an ASD or POP classification, and a sufficient amount of DNA available for methylation measurements.

A complete description of the SSC, which enrolled and collected biospecimens from children and adolescents aged 4-18 years, can be found elsewhere [37]. Briefly, a geneticist and a clinical psychologist were appointed as co-principal investigators at each site. Probands were evaluated with a battery of diagnostic measures, including the Autism Diagnostic Interview – Revised (ADI-R) [34] and the Autism Diagnostic Observation Schedule (ADOS) [31]. Other instruments provided additional measures of the core features of autism, as well as of intellectual ability (verbal and nonverbal), adaptive behavior, emotional and behavior problems, motor function, and language. A description of instruments employed can be found at https://sfari.org/ssc-instruments. A comprehensive family medical history was obtained that included the proband’s prenatal and perinatal history, developmental milestones, immunizations, medications, dietary supplements, and common behavioral treatments. Emphasis was placed on common “comorbidities” including gastrointestinal complaints, sleep irregularities, and seizures. In addition, questions were asked about genetic, autoimmune, and psychiatric disorders in members of the extended family. Probands were excluded who were younger than 4 years of age or older than 18. Probands were also excluded for conditions that might compromise the validity of diagnostic instruments, such as nonverbal mental age below 18 months, severe neurological deficits, birth trauma, perinatal complications, or genetic evidence of fragile X or Down syndromes. A complete description of exclusion/inclusion criteria can be found at http://sfari.org. Measures of adaptive function, behavior-emotional problems, and symptoms of autism were examined in parents and siblings as well as probands. Thus, the SSC represents a unique, well-described sample of able children and adolescents with relatively severe ASD, as indicated by ADI-R and ADOS Calibrated Severity Scores [38].

Reliability of Data: to maximize the consistency of clinical observations across sites, each clinician was trained in administration of the ADOS and ADI-R to achieve research reliability as judged by expert clinicians. Most clinicians who had not previously received research training required 4–6 months of practice. Videotapes of interviews were exchanged to ensure that reliability requirements were met and maintained throughout the study. Error rates were very low, averaging less than 0.50 errors/1000 data points. Most errors could be corrected immediately, resulting in an unusually clean data set for a multisite study of this size. During each visit, a blood sample was collected from each study participant and DNA was extracted from blood cells, while plasma was stored for future use.

### DNA methylation data quality control (QC) and processing

For the SEED samples, genomic DNA was isolated from 980 whole blood samples using the QIAsymphony midi kit (Qiagen). For each sample, 500 ng of DNA was bisulfite treated using the 96-well EZ DNA methylation kit (Zymo Research). Samples were randomized within and across plates, and across two main processing dates to minimize batch effects, and run on the Illumina HumanMethylation450 BeadChip. Background correction and dye-bias equalization was performed using the function *preprocessNoob()* [39, 40] in the *minfi* R package [41]. We included 12 cross-plate duplicates for quality control purposes; pairwise correlation metrics for the duplicate samples ranged from 0.990 to 0.997 with a mean correlation value equal to 0.995. Samples were removed if they had low overall intensity (median unmethylated or methylated signal < 11) or had a detection p-value > 0.01 in more than 1% of probes (N = 7), or if reported sex did not match predicted sex generated via the *minfi* function *getSex()* (N = 3). Probes were removed if they had a detection p-value > 0.01 in more than 10% of samples (n = 702) and then if they had been previously identified as being ambiguously mapped [42] (n = 29,146). Following QC, the analytic data included DNAm for 455,664 sites on 970 samples. We further removed 2 samples who were missing a final outcome classification, leaving a total of 453 cases and 515 controls used for association analyses.

For the SSC samples, five hundred nanograms of human genomic DNA was sodium bisulfite-treated for cytosine to thymine conversion using the EZ DNA Methylation Gold kit (Zymo Research). A total of 728 samples (from 364 families) were randomized within and across plates to minimize batch effects, and run on the Illumina HumanMethylation450 BeadChip. Additional details have been previously described [43]. Similar quality control procedures as used for the SEED samples were used for the SSC samples. After background correction and dye-bias equalization, samples were removed for low overall intensity (median unmethylated or methylated signal < 11) or for detection p-value > 0.01 in more than 1% of probes (N = 42). Probes were removed if they had a detection p-value > 0.01 in more than 10% of samples (n = 483) and then if they had been previously identified as being ambiguously mapped (n = 29,213). These steps resulted in an analytic data set with 455,816 sites on 686 samples, consisting of 343 proband-sibling pairs.

Finally, for all SEED and SSC samples we estimated cell type proportions for 6 different cell types (granulocytes, monocytes, CD4 T cells, CD8 T cells, B cells, and natural killer cells) using the *estimateCellCounts()* function in the *minfi* R package. Estimation incorporated reference data from 60 samples generated from 6 healthy adult men [44].

### Genotype data quality control and processing

Whole genome genotyping data was available for 943 of the 970 SEED 1 samples which passed DNAm quality control steps. After genotype measurement using the Illumina HumanOmni1-Quad BeadChip, standard quality control measures were applied, including removal of samples with < 95% SNP call rate, sex discrepancies, relatedness (Pi-hat > 0.2), or excess hetero- or homozygosity, and removal of markers with < 98.5% call rate, or that were monomorphic. Phasing was performed using SHAPEIT [45] followed by SNP imputation via the IMPUTE2 software [46], using 1000 Genomes Project samples as reference. Principle components to account for ancestry were determined via the EigenStrat program [47].

### Epigenome-wide Association testing and meta-analysis

For the SEED data, we used linear regression modeling of the M-value (the ratio of methylated to total signal determined at every probe in every sample) [48] as a dependent variable and ASD status, sex, and surrogate variables (SVs) (described below) as independent variables. We implemented this model using the *lmFit()* function in the *limma* R package [49], separately for each of the DNAm probes that passed QC. For the SSC data, we implemented a generalized estimating equation (GEE) model using the *gee()* function in the *gee* R package [50] to account for the correlation inherent to the familial structure in the data. We used a fixed correlation structure of 0.5 for each sibling pair, and regressed M-value onto ASD status, sex and SVs.

To account for sources of technical and biological variability in our association analyses, we estimated surrogate variables (SVs) [51] in the cleaned SEED and SSC dataset to include as covariates in our downstream analyses. SVs have been shown to capture and adjust for differences related to batch effects and cell type proportions across samples in a wide variety of simulated settings [52], and to remove the effects of unwanted sources of technical and biological variation [51]. In order to explicitly address the strong confounding effect of sex resulting from the high degree of male bias in ASD diagnosis, we removed sex chromosomes, where DNAm values strongly correlate to sex, before SV estimation, and included sex along with ASD status in the model used for SV estimation. We then used a data-driven procedure individually in the SEED and SSC data to select the number of SVs to include in the association models.

First, to examine the relationship between each SV and known sources of technical and biological variation, we estimated the association between each estimated SV and cell type composition, principal components of genetic ancestry, and processing batch. We then generated a visual representation of the degree of association with these variables using a heat map (**Additional File 1: Figures S1a, S2a**).

We next examined the influence of iterative inclusion of SVs as adjustment variables in our association regression models. To do this, we first ran a case-control association model with adjustment for the strongest estimated SV [51], then progressively included the next strongest SV in the analysis and continued this procedure until all estimated SVs were included. For each model, we recorded the inflation factor, or lambda, calculated via the *estlambda()* function from the *GenABEL* R package [53], and visualized the relationship between number of SVs adjusted for and lambda values (**Additional File 1: Figures S1b, S2b**). We chose the number of SVs to include in the model by considering both the number of SVs at which the estimated lambda values began to plateau and where the known potential confounders appeared to be captured by one or more SVs. We chose to include 19 SVs in the SEED association analysis and 14 in the SSC analysis.

After completing each association analysis, we then performed a meta-analysis using the METAL software [54] on the 445,068 probes that were present in both the SEED and SSC cleaned datasets. Our approach weighted individual study effects by sample size and also took into account the direction of effect. We also computed the false discovery rate (FDR) using the Benjamini-Hochberg method [55]. We also determined statistical power for this meta-analysis *a priori* using an estimation method specifically designed for epigenome-wide association studies [56].

### Comparison of blood EWAS hits to brain-based DNAm

We sought to compare the consistency of top EWAS results from the blood-based meta-analysis to our previous analysis of post-mortem brain samples from ASD cases and controls [19]. These data consist of DNAm from three brain regions: cerebellum, prefrontal cortex, and temporal cortex. For the CpG sites reaching suggestive levels of significance (p < 1×10^−4^) in the meta-analysis, we computed mean differences between cases and controls in each of these 3 brain regions. We then computed Pearson correlations and quadrant count ratios between the blood effect sizes and 3 lists of brain effect sizes. We computed quadrant count ratios as the sum of concordant effect sizes (both positive or both negative) minus the sum of discordant effect sizes, all divided by the total number of effect sizes being compared.

### Methylation quantitative trait loci (meQTL) query and meQTL target enrichment test

We were interested in exploring the propensity of CpG sites that reached a level of suggestive significance in the EWAS meta-analysis to be significantly associated with nearby SNPs. We used joint DNAm and genotype data to define SNPs associated DNAm, sometimes referred to as “methylation quantitative trait loci (meQTLs)”, and the CpG sites under genetic control, or “meQTL targets”. We then tested for enrichment of meQTL targets in the top ranked CpG sites from the meta-analysis.

In lieu of applying our SV selection method (see ‘Association testing and meta-analysis’) to every SNP-CpG association test in the meQTL query, we conducted separate meQTL queries in each processing batch of the SEED data (N_Batch1_ = 606; N_Batch2_ = 362). In each batch, we first used a data-driven procedure we have described in detail previously [57] to select three key parameters for the meQTL query: the SNP minor allele frequency threshold for inclusion, the CpG variability threshold for inclusion, and the maximum distance between SNP and CpG site to be considered for analysis. Briefly, this procedure selects parameters to ensure 80% power to detect a 5% DNAm difference with each addition of the minor allele, at a Bonferroni-defined significance threshold. We then performed the meQTL query in each batch using the *MatrixEQTL* R package, adjusting for sex, the first 5 principle components to account for genetic ancestry, and the first 2 principle components derived from the cell composition estimates. We then defined SNP-CpG association pairs based on results that can gain 100% power in the parameter survey (“permissive”), 90% power (“intermediate”), and 80% power (“stringent”). If a SNP-CpG association pair was significant at a designated threshold in each batch, the CpG site was labeled a meQTL target under that threshold for the downstream enrichment analysis.

We tested for enrichment of meQTL targets in ASD-associated CpG sites. We examined this using two ASD EWAS meta-analysis p-value thresholds (p<1×10^−3^ and p<1×10^−4^) and the three meQTL p-value thresholds. In each enrichment test, we accounted for the two main features of CpG sites likely to affect results: the degree of variability in DNAm at that CpG and the number of SNPs in the boundary considered. To do this, we binned each CpG site by decile according to these factors. For each EWAS/meQTL threshold scenario, we compared the proportion of meQTL targets among ASD-related CpGs to a null distribution of randomly selected CpGs, equal in count to the number of ASD-associated sites, matched on the same variability and nearby SNP decile. We defined a fold enrichment statistic as the count of meQTL targets in the ASD-associated CpGs divided by the mean proportion of meQTL targets from the null set, and an enrichment p-value as the number of null CpG sets with a count of meQTL targets that was equal to or exceeded the count in the ASD-associated CpG list.

## Results

### ASD EWAS meta-analysis in blood

We performed a meta-analysis over the 445,068 probes that were present in both the SEED (**Additional File 2: Table S1**) and SSC (**Additional File 2: Table S2**) cleaned datasets. Figure 1A shows the range of p-values and effect sizes detected in our meta-analysis. No CpG sites reached a Bonferroni level of significance, and effect sizes were modest (1.12×10^−7^; Figure 1A). The genomic inflation factor (λ) was 1.03, with a slight separation from expectation at the tail (Figure 1B). A total of 48 CpG sites met or exceeded a p-value < 1×10^−4^ and 7 CpG sites (Table 1) reached a significance level of p < 1×10^−5^. We have provided a full list of summary statistics for both SEED and SSC for all 445,068 probes (Additional File 3: **Table S3**). Based on our analytic sample size, we had 80% power to detect a 3.8% DNAm difference between cases and controls at a Bonferonni level of significance.

**Table 1.**
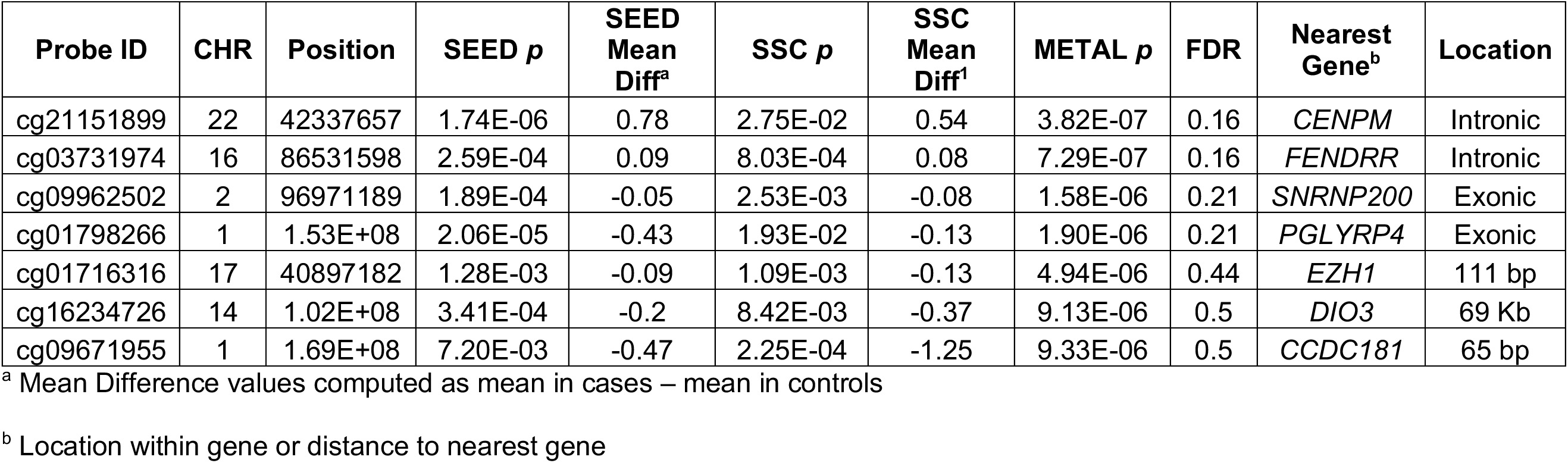
CpG sites identified from meta-analysis as being suggestively associated (1×10^−5^) with ASD.

**Figure 1.**
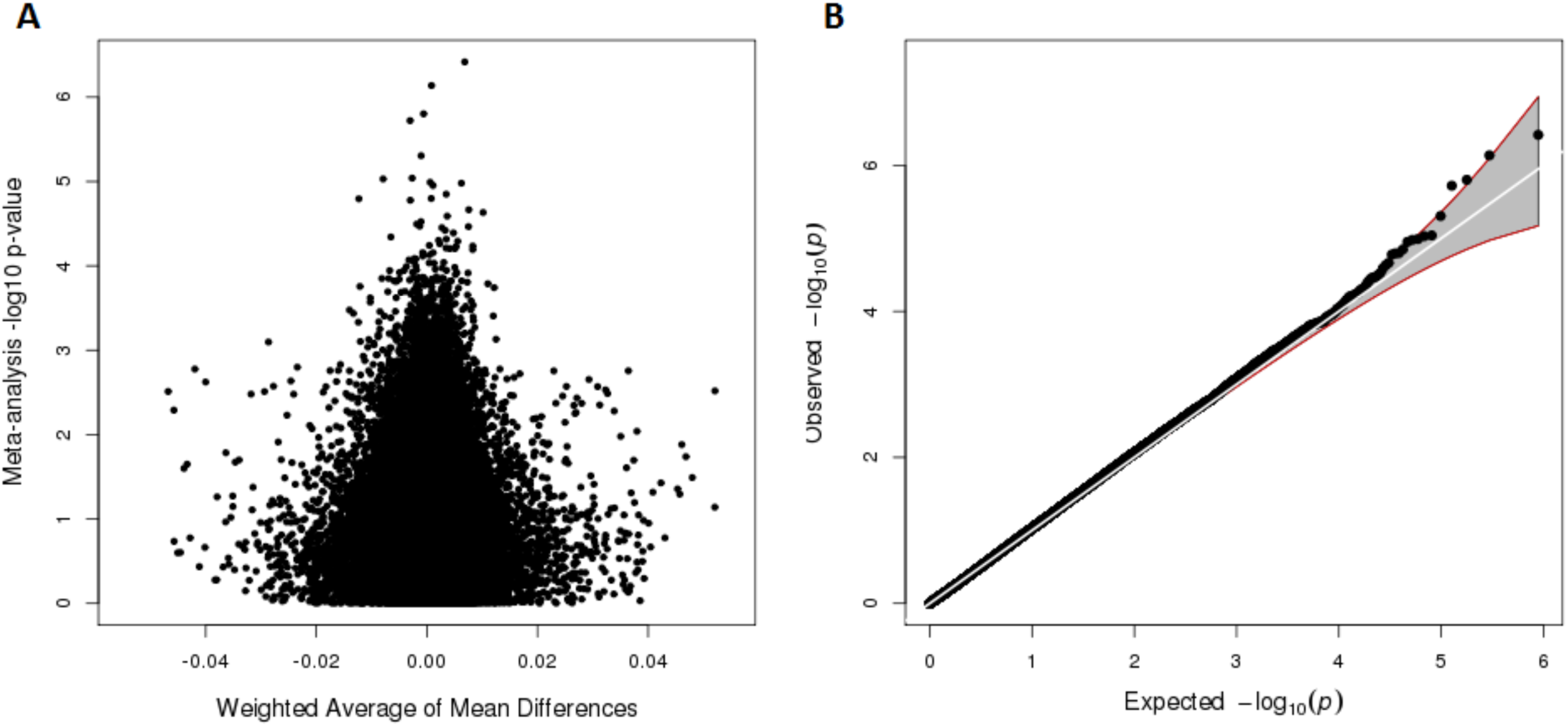
Meta-analysis results of epigenome-wide association analysis for ASD in peripheral blood in SEED in SSC samples. Panel A) Volcano plot depicting meta-analysis p-value (log base 10 scale) on y-axis and average of mean difference values in SEED and SSC samples weighted by sample size on x-axis. Panel B) Quantile-quantile plot (λ = 1.03).

### Consistency of blood EWAS hits in brain

We considered the consistency of signal for the 48 blood-based CpGs with suggestive significance (p-value < 1×10^−4^), among results from three different brain regions with data available from our previous analysis of post-mortem brain samples and ASD [19]. The cerebellum exhibited a moderate degree of concordance in effect size and direction (r = 0.31; QCR = 0.33); although prefrontal cortex (r = 0.02; QCR = 0.125) and temporal cortex (r = −0.10; QCR = −0.125) showed only minimal concordance (Additional File 4: Table S4, Additional File 5: Figure S3). When considering the 7 CpG sites with more stringent blood-based p-values < 1×10^−5^, the direction of effect was consistent for at least 5 of these 7 in all three brain region results, with typically larger effect sizes (Table 2). The CpG site with the largest effect size in blood (cg09671955) displayed consistent, and larger, effect sizes in all three brain regions (Table 2, Additional File 4: Table S4).

**Table 2.** Suggestively associated (p < 1×10^−5^) CpGs sites in peripheral blood and their corresponding effect sizes in three brain regions.

| Probe ID | CHR | Position | SEED Mean Diff<sup>a</sup> | SSC Mean Diff<sup>a</sup> | Weighted Average Mean Diff<sup>b</sup> | PFC<sup>a,c</sup> | TC<sup>a,d</sup> | CER<sup>a,e</sup> |
| --- | --- | --- | --- | --- | --- | --- | --- | --- |
| cg21151899 | 22 | 42337657 | 0.78 | 0.54 | 0.68 | 1.60 | 4.07 | 1.64 |
| cg03731974 | 16 | 86531598 | 0.09 | 0.08 | 0.08 | 0.20 | -7.18 | 0.74 |
| cg09962502 | 2 | 96971189 | -0.05 | -0.08 | -0.06 | -2.50 | -0.43 | -1.34 |
| cg01798266 | 1 | 1.53E+08 | -0.43 | -0.13 | -0.31 | -2.70 | -0.91 | 0.87 |
| cg01716316 | 17 | 40897182 | -0.09 | -0.13 | -0.1 | -0.30 | -0.02 | 0.39 |
| cg16234726 | 14 | 1.02E+08 | -0.2 | -0.37 | -0.27 | 0.50 | -0.3 | -0.03 |
| cg09671955 | 1 | 1.69E+08 | -0.47 | -1.25 | -0.79 | -1.40 | -1.38 | -5.11 |

<sup>a</sup> Mean Difference values computed as mean in cases – mean in controls
<sup>b</sup> Average of SEED and SSC mean difference values weighted by sample size ( $N_{\text{SEED}} = 968$ , $N_{\text{SSC}} = 686$ ).
<sup>c</sup> Prefrontal cortex data from Ladd-Acosta et al. [19]
<sup>d</sup> Temporal cortex data from Ladd-Acosta et al. [19]
<sup>e</sup> Cerebellum data from Ladd-Acosta et al. [19]

### meQTL target enrichment test

When considering all CpGs associated with ASD at a liberal p < 1×10^−3^ EWAS threshold, we found significant meQTL target enrichment (p_enrichment_ = 0.041) (Table 3). All other combinations of EWAS and meQTL p-values displayed suggestive levels of significance (0.089 ≤ p_enrichment_ ≤ 0.243) and a consistent direction of effect towards enrichment. Also, tests conducted for CpGs meeting the more stringent EWAS p-value threshold (1×10^−4^) displayed a consistently greater effect size than their corresponding tests from the more liberal EWAS threshold.

**Table 3.** Enrichment statistics for meQTL targets in ASD-related CpG sites.

|  |  | meQTL p-value threshold<sup>b</sup> |  |  |
| --- | --- | --- | --- | --- |
|  |  | Permissive<sup>c</sup> | Intermediate<sup>d</sup> | Stringent<sup>e</sup> |
| **ASD EWAS p-value<sup>a</sup>** | **1x10<sup>-3</sup>** | 1.20 (0.041) | 1.11 (0.218) | 1.11 (0.243) |
|  | **1x10<sup>-4</sup>** | 1.46 (0.134) | 1.71 (0.089) | 1.50 (0.205) |

Enrichment fold statistics and p-values based on 1,000 permutations are reported.
<sup>a</sup>ASD to DNAm association p-value defined from meta-analysis
<sup>b</sup>SNP to CpG association p-value thresholds
<sup>c</sup>p-values in each methylation processing batch that allowed for 100% power to detect a 5% methylation difference with each addition of minor allele (see Methods).
<sup>d</sup>p-values in each methylation processing batch that allowed for 90% power to detect same methylation difference
<sup>e</sup>p-values in each methylation processing batch that allowed for 80% power to detect same methylation difference.

## Discussion

We provide the largest study to date investigating the relationship between ASD and DNAm. A case-control meta-analysis of peripheral blood samples from the Study to Explore Early Development and the Simons Simplex Collection revealed that none of the 455,068 CpG sites investigated were associated with ASD at a genome-wide significant threshold. However, 48 CpG sites reached suggestive significance levels at p < 1×10^−4^, including 7 CpGs at p < 1×10^−5^. Associations with ASD at these sites display moderate concordance with post-mortem brain sample results from a previous study and display evidence for enrichment of SNP-controlled CpG sites, or meQTL targets.

Given the potential involvement of epigenetic mechanisms in ASD [4–12], and the availability of blood samples from existing studies, this was an important project to pursue. In contrast to our null findings in blood samples, previous work using brain samples has shown specific DNAm to be associated with ASD [15, 19, 20]. The difference in tissue type might explain the inconsistent results across these studies. Nonetheless, a previous study of ASD and DNAm in peripheral blood [24] using the earlier 27K Illumina array reported numerous differentially methylated sites. Our study, despite a much higher sample size, did not observe associations at these same sites. However, their reported differentially methylated sites were based on a ranking that integrated both degree of statistical significance and effect size; none of their single CpG associations achieved statistical significance at a Bonferroni correction level. Also, they did not explicitly account for potential confounding by cell type proportions or address potential batch effects in their analytic pipeline, as this was not yet commonplace in EWAS pipelines at the time of their report. We used a rigorous, data-driven method to account for these factors and control the genome-wide Type I error rate. It is also possible that true differentially methylated positions for ASD exist in blood, but have smaller effects sizes than we were powered to detect. Recent EWAS discoveries have indeed found replicable very small effect sizes, particularly in environmental health [58].

Importantly, our study is based on a case-control design using samples post-onset of ASD, rather than biosamples taken early in development. Thus, our ability to examine ASD etiologic mechanisms is limited. However, there are many recent examples where blood-based epigenetic work in brain-based disorders can be useful, despite such a limitation [57, 59–63]. For example, DNAm under genetic control (meQTL targets) can inform genetic associations observed for ASD. Further, consideration of aggregated sets of CpGs associated with ASD, rather than single sites, can elucidate pathways of interest [57]. In our own analysis, the most differential blood-based CpGs had consistent effect sizes and directions, although weaker, among brain-based results, particularly for cerebellum. Blood-based CpGs were also moderately enriched for meQTL targets. These results suggest that blood DNAm can be reflective of DNAm in affected tissues, and suggest genetic control of DNAm as a mechanism for this occurrence, at least in an ASD context. More precise evidence is needed, but given the easy accessibility of blood for DNAm measurements versus brain [61], the utility of blood-based DNAm research in ASD is worthy of additional consideration.

In summary, our work is the largest study of DNAm and ASD to date. Our results point to the need for even larger studies to take place in the future, and argue for an ongoing investigation of the role of genetic factors in contributing to DNAm differences in ASD. To this end, we have provided our full summary statistics and meta-analysis results. The need for greater sample sizes mimics the initial stages of genetic variation discovery in ASD, for which large mega-analyses are starting to pay dividends [64].

## List of abbreviations

**ASD**: autism spectrum disorder; **DNAm**: DNA methylation; **EWAS**: epigenome-wide association study; **meQTL**: methylation quantitative trait loci; **SEED**: Study to Explore Early Development; **SNP**: single nucleotide polymorphism; **SSC**: Simons Simplex Collection; **SV**: surrogate variable

## Declarations

### Ethics approval and consent to participate

This study was approved by the institutional review boards at each SEED site: SEED 1 recruitment was approved by the IRBs of each recruitment site: Institutional Review Board (IRB)-C, CDC Human Research Protection Office; Kaiser Foundation Research Institute (KFRI) Kaiser Permanente Northern California IRB, Colorado Multiple IRB, Emory University IRB, Georgia Department of Public Health IRB, Maryland Department of Health and Mental Hygiene IRB, Johns Hopkins Bloomberg School of Public Health Review Board, University of North Carolina IRB and Office of Human Research Ethics, IRB of The Children's Hospital of Philadelphia, and IRB of the University of Pennsylvania. All enrolled families provided written consent for participation. This methylation substudy was approved as an amendment of the Johns Hopkins Institutional Review Board (IRB) approval. For participants from the Simons simplex collection, parents consented and children assented as required by each local institutional review board, which included a coalition of clinics located at Michigan, Yale, Emory, Columbia, Vanderbilt, McGill Washington, and Harvard Universities (Children’s Hospital of Boston), and at the Universities of Washington, Illinois (Chicago), Missouri, UCLA, and the Baylor College of Medicine. To protect the privacy of participants, Global Unique Identifiers (GUIDs) were constructed from personal information using an algorithm devised in were collaboration with scientists at the NIH [65]. Each clinic retained personal identifiers on site and transmitted de-identified GUIDs to a central database, as described previously [37].

### Consent for publication

Not applicable.

### Availability of data and material

The SEED I data generated and analyzed for this study are not publicly available due to lack of explicit consent for such sharing in the written informed consents for SEED sites, according to the CDC IRB that governs the SEED network. The SEED data generated and analyzed for this study are not publicly available due to confidentiality and informed consent requirements. We have submitted the raw 450K methylation data from the SSC samples to the National Database for Autism Research (NDAR; ndar.nih.gov), which can be found under collection #300.

### Authors’ contributions

MDF and CL-A conceived the study. GCW, LAS, DES, LAC, MDF, and CJN led participation and sample recruitment for SEED. STW, RSA and PC generated the methylation data and performed initial analysis on all the SSC samples. CL-A and APF supervised methylation data collection. SVA performed quality control and preprocessing for the SEED and SSC methylation data. BS and CL-A performed quality control for the SEED genotype data. SVA performed all analyses, supervised by MDF and CL-A. CL-A, MDF, and APF obtained funding for the DNA methylation measurements in SEED. SVA, MDF, and CL-A wrote the manuscript. All authors contributed to interpretation of results and edited and reviewed the manuscript.

## Additional File

**Additional File 1 – Figures S1-S2. Depiction of surrogate variable selection process for SEED (S1) and SSC (S2).** Panel A: Heatmap indicating degree of association with known potential technical variables or confounders with estimated surrogate variables. Panel B: Inflation factor (lambda) calculated for progressively including surrogate variables in association models. The number of surrogate variables to include in the ultimate association testing model was to determine to be that which properly controlled the inflation factor and adequately captured known technical variables or confounders. See Methods for additional explanation.

**Additional File 2 – Tables S1-S2. Demographic characteristics for samples in the SEED (S1) and SSC (S2) datasets.**

**Additional File 3 – Table S3. Full summary statistics and meta-analysis results for all 445,608 CpG sites that were present in both the cleaned SEED and SSC datasets.**

**Additional File 4 – Table S4. Concordance between suggestively associated (p < 1×10^−4^) CpG sites in peripheral blood and their corresponding effect sizes in three brain regions**

**Additional File 5 – Figure S3. Quadrant plots depicting concordance in effect sizes between suggestively associated (p < 1×10^−4^) CpG sites in peripheral blood and three brain regions.** A) Prefrontal cortex B) Temporal Cortex C) Cerebellum. Points in red indicate those sites with p < 1×10^−5^ in peripheral blood.

